# Analysis of a European general wildlife health surveillance program: chances, challenges and recommendations

**DOI:** 10.1101/2023.07.13.548813

**Authors:** Elisabeth Heiderich, Saskia Keller, Mirjam Pewsner, Francesco Carlo Origgi, Samoa Zürcher-Giovannini, Stéphanie Borel, Iris Marti, Patrick Scherrer, Simone Roberto Rolando Pisano, Brian Friker, Irene Adrian-Kalchhauser, Marie-Pierre Ryser-Degiorgis

## Abstract

In the context of a One Health perspective general wildlife health surveillance (GWHS) gains importance worldwide, as pathogen transmission among wildlife, domestic animals and humans raises health, conservation and economic concerns. However, GWHS programs operate in the face of legal, geographical, financial, or administrative conditions. The present study uses a multi-tiered approach to understand the current characteristics, strengths and gaps of a European GWHS that operates in a fragmented legislative and multi-stakeholder environment. The aim is to support the implementation or improvement of other GWHS systems by managers, surveillance experts, and administrations.

To assess the current state of wildlife health investigations and trends within the GWHS, we analyzed 20 years of wildlife diagnostic data, conducted an online survey and phone interviews with official field partners (hunting administrators, game wardens and hunters), and performed a time-per-task analysis.

Firstly, we found that infectious disease dynamics, the level of public awareness for specific diseases, research activities and increasing population sizes of in depth-monitored protected species, together with biogeographical and political boundaries all impacted case numbers and can present unexpected challenges to a GWHS. Secondly, we found that even a seemingly comprehensive GWHS can feature pronounced information gaps, with underrepresentation of common or easily recognizable diseases, blind spots in non-hunted species and only a fraction of discovered carcasses being submitted. Thirdly, we found that substantial amounts of wildlife health data may be available at local hunting administrations or disease specialist centers, but outside the reach of the GWHS and its processes.

In conclusion, we recommend that fragmented and federalist GWHS programs like the one addressed require a central, consistent and accessible collection of wildlife health data. Also, considering the growing role of citizen observers in environmental research, we recommend using online reporting systems to harness decentrally available information and fill wildlife health information gaps.

## Introduction

### Wildlife health surveillance (WHS) is an integral aspect of One Health inspired national health management strategies [1,2]

The repeated emergence of human and livestock diseases from wildlife origins and the increasing risk of pathogen transmission from wild animals to humans and domestic animals associated with patterns of global change, underline the importance of WHS [3–5]. Emerging diseases in wildlife also contribute to biodiversity loss [6]. For these reasons, WHS constitutes an essential component of national and organizational preparedness strategies [1,7–9]. Accordingly, wildlife health has become a priority for the World Organization for Animal Health (WOAH, formerly OIE) not only in the wake of the COVID-19 pandemic [10]).

### General wildlife health surveillance (GWHS) is a major component of WHS programs

The term stands for a passive system of opportunistic carcass collection that encompasses a wide range of species and diseases [8,11]. It is often complemented by targeted disease- or species-specific focus programs (active or targeted surveillance), and complemented with population monitoring data, to assess epidemiological dynamics, freedom of disease and intervention outcomes in an integrated wildlife monitoring (IWM) program [11].

### Features characterizing effective GWHS have been described previously

The aim of a national GWHS program should be the identification, effective communication and management of risks to or from the country’s wildlife populations [2]. For example, GWHS programs need to assure early detection of emerging pathogens, which is a prerequisite for rapid reactions towards disease control [12–14]. They should also generate appropriate knowledge to improve the effectiveness of wildlife policies and systems [2]. As wildlife diseases do not stop at country borders, harmonization and centralization of information at an international level is of high importance to increase efficiency of national programs [11].

### In practice, however, GWHS faces several challenges

These include aspects related to sample availability, but also to expertise, legal frameworks, and resources. Firstly, GWHS depends on spontaneous reports on wildlife morbidity and mortality by field partners (state game wardens, hunters, wildlife biologists) and from the public. In practice, this means that dead or diseased wildlife must be found, reported, sent for pathological examination, in the shortest possible time to prevent that carcass decomposition may mask the actual pathological lesions, for effective surveillance. Therefore, GWHS can be biased against elusive, common, or non-hunted species. Secondly, GWHS deals with species uncommon to most veterinarians, and often face a lack of validated diagnostic tests, baseline data, harmonized procedures, population data, and expertise [14,15]. Finally, the benefit of a wildlife health surveillance program to society is difficult to express in metrics, and the fact that wildlife is a public good limits the resources available to GWHS [1,2,16].

### GWHS often operates in the face of non-negotiable frame conditions

These include (a) the reactivity of the system to natural developments (e.g., disease outbreaks or changes in population density), (b) the reactivity of the system to societal developments (e.g., new laws, mandates, increased attention on biodiversity or One Health), (c) the dependencies and perspectives of submitters and funding stakeholders, their needs, limitations, and expertise, (d) the de-facto entanglement and co-dependencies among general surveillance, targeted programs, and research, (e) historically grown and legislatively set boundaries and practices, and (f) potential information gaps associated with all of the above. GWHS is expected to operate and provide valuable insights despite these limitations.

### The GWHS program analyzed here has a comprehensive scope and was started around 1950 in Switzerland [7,8]

Switzerland is a small federal state (41’285 km^2^), but highly heterogeneous in terms of landscape, climate, habitat, human population density, language, political organization and wildlife management. Twenty-six variably sized, biogeographically distinct, and largely independent political subunits (“cantons”, Fig 1) are further subdivided in wildlife management districts of different sizes [17,18]. The adjacent Principality of Liechtenstein (160 km^2^) is, for the purpose of veterinary services, affiliated with the Swiss systems.

**Fig 1.**
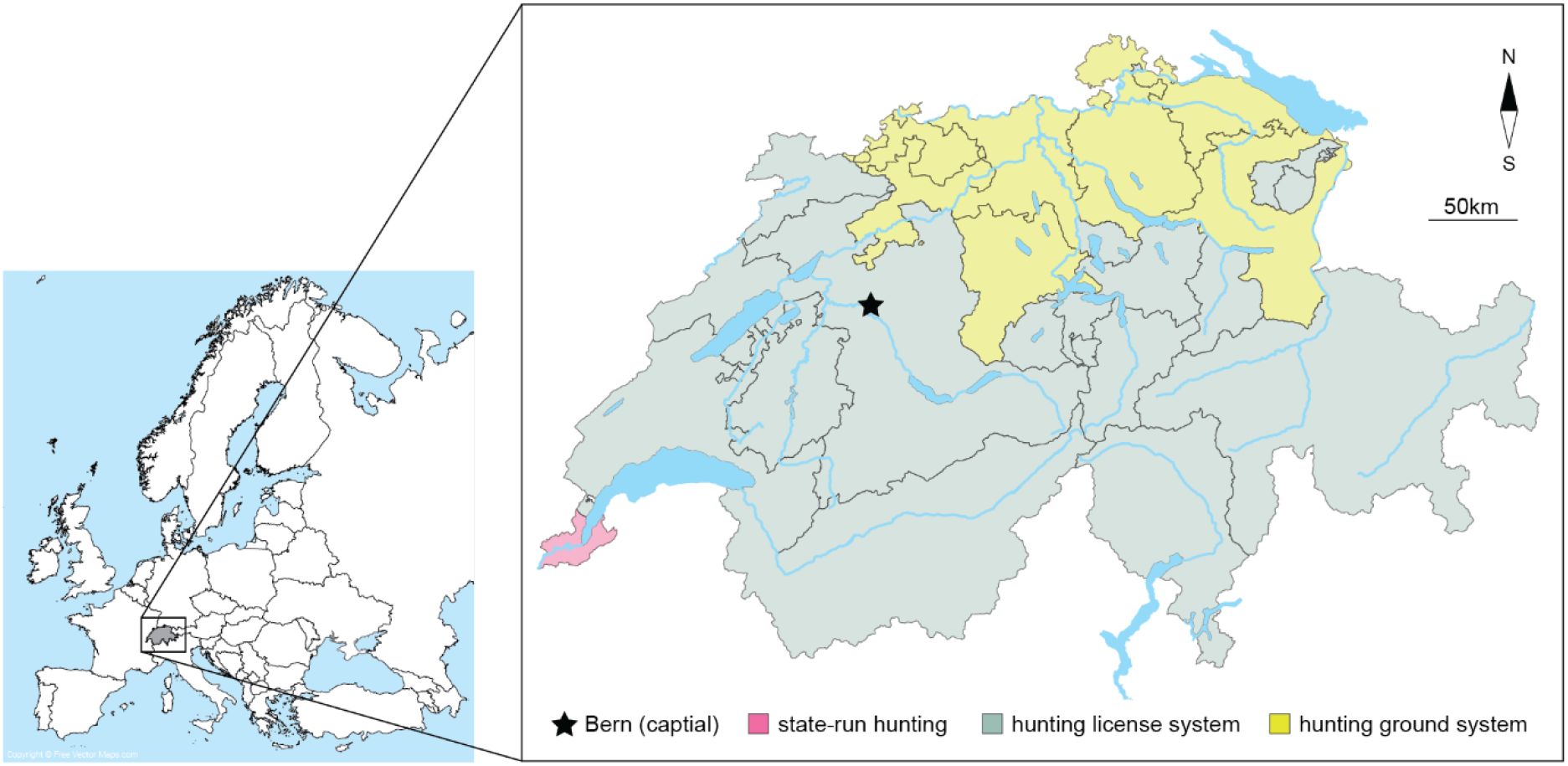
Study setting. The Swiss general wildlife health surveillance system operates in a geographically limited, but administratively fragmented setting. Switzerland is a federative state comprising 26 states (“cantons”) which feature three distinct hunting regimes and vastly different terrains and habitats, from agricultural to high alpine. The area also features major biogeographic divides, i.e., four river basins and the main crest of the alps. The Institute for Fish and Wildlife Health is located in the capital city Bern.

### The Swiss terrestrial wildlife management system is legally, geographically and administratively fragmented

The Swiss legal system defines wildlife as *res nullius*, which means they are not the subject of private property but governmentally managed. It is important to understand that wildlife health is regulated by three different laws: the Federal Act on Hunting and the Protection of Wild Mammals and Birds (hunting law) [19], the Federal Act on Animal Diseases (animal disease law) [20] and the Federal Act on the Protection of Nature and Cultural Heritage (nature protection law) [21]. The cantons are responsible for the legal execution of these laws. Each canton features one of three hunting regimes: (1) the hunting license system (16 cantons and the Principality of Liechtenstein), (2) the hunting ground system (nine cantons), and (3) state-run hunting (one canton) (Fig 1). Officially appointed professional state game wardens are present in all cantons with the license system and in the state-run hunting system [22]. In cantons with hunting grounds, these tasks are partly performed by voluntary game wardens. The health surveillance of species not covered in the hunting law is subject to the nature protection law, which is primarily concerned with biodiversity and species protection and conservation and coordinated by the Federal Office for the Environment (FOEN; different division than hunting). The animal disease law focuses on zoonotic diseases or pathogens with high risk of spill-over to domestic and farmed animals, and to any species these pathogens may pertain, and is coordinated and enforced on the federal level by the Federal Food Safety and Veterinary Office (FSVO).

### The Swiss GWHS relies on a single diagnostic institute

Currently, GWHS for terrestrial animals is conducted at the University of Bern, Veterinary Faculty, Institute for Fish and Wildlife Health (FIWI), hereafter referred to as diagnostic institute, on a mandate of the FOEN and the FSVO. Three academic, three cantonal and one private veterinary laboratory in Switzerland carry out wildlife necropsies. However, only the diagnostic institute is specialized in wildlife and postmortem disease diagnostics.

### The diagnostic institute has three mandates in addition to GWHS

The GWHS covers (a) common free-ranging, middle sized to large wild mammals and birds including those which can be legally hunted (species covered by the hunting law) [19]; (b) eulipotyphla, bats, small rodents, reptiles and amphibians that are not hunted (species covered by the nature protection law). In addition, the diagnostic institute (1) carries out an in-depth health monitoring mandate for protected wild mammals, including the Eurasian lynx (*Lynx lynx*), gray wolf (*Canis lupus*), golden jackal (*Canis aureus*), brown bear (*Ursus arctos*) as well as, until 2019, the Eurasian beaver (*Castor fiber*); (2) expert reports on predation, with the aim to identify the predator species for farmed or domestic animals and (3) postmortem investigations in farmed deer species.

### Additional institutions execute targeted surveillance programs for specific pathogens

Targeted surveillance programs for pathogens that also affect wildlife are carried out by a variety of institutions, e.g., National Reference Centre for Poultry and Rabbit Diseases, University of Zurich (avian influenza), Institute for Virology and Immunology, joint institute University of Bern and FSVO (African swine fever), Department for Veterinary Bacteriology, University of Zurich (tuberculosis), and Institute for Parasitology, University of Bern (trichinellosis). The host institutions of targeted programs are usually specialized on the respective pathogen group (bacteriology, virology, parasitology, etc.) or host taxon (e.g., birds). However, they often have a focus on domestic/farmed animals rather than wildlife. The diagnostic institute submits carcasses or samples to these programs on occasion for the targeted surveillance of reportable diseases, to prove freedom of reportable diseases or based on pathological signs and suspicion (bats and foxes for rabies, lynx and wolves for trichinellosis, brown hare for tularemia, etc.).

### The GWHS has characteristics of an academic setting

The location at the University of Bern comes with certain hallmarks of academic institutions: a dependency on multiple intra- and extra-mural funding sources; the expectation to use submitted cases for teaching; the involvement of fast turn-over trainees and the need for supervision in diagnostics; the availability of in-depth expertise on various diseases and pathogen groups from in-house specialized research groups; the ability, but also the expectation to turn unexpected findings into research projects; the expectation of long-term employees to be able to participate in activities other than diagnostic (own research, teaching, board activities).

### The GWHS depends on multiple funding sources

Salaries of diagnostics employees are largely funded by the two aforementioned federal offices. The university contributes infrastructure and administration, the salary of the institute head and two to three research trainee positions. Submitters are usually not charged for wildlife postmortem examination including further ancillary testing, except for toxicological investigations and postmortem examinations of farmed deer.

### This paper provides an unredacted insight into a European GWHS system operating within real-world constraints

Increased numbers of case submissions for postmortem investigation despite constant funding over the past twenty years motivated this study. Analyses include (1) a detailed breakdown of cases from 2002-2019 including causes of death, submitter types, geographical origin and comparison to other national data sources; (2) qualitative interviews and online surveys to capture the perspective and needs of submitters, program partners and funders; and (3) analyses of aspects that disproportionally affect time per case. Aim of these analyses was to identify unmet needs and data gaps, suggest measures to make better use of the limited financial resources, and identify future opportunities for GWHS.

### With this paper we want to support the implementation or improvement of other real-world GWHS systems by managers, surveillance experts, and administration

We provide an example on how to analyze a running system and create awareness for the multidimensional space of needs and framework conditions in which GWHS programs operate. We develop a picture of GWHS strategic management as a fluid, interactive, and transdisciplinary process of continuously co-creating and balancing a non-perfect multi-stakeholder system. Finally, we come to conclusions and provide several data-supported recommendations on how to improve, or to build, GWHS in the face of the described settings.

## Material and Methods

### Retrospective case analysis

#### To assess the current state of wildlife investigations and trends within the GWHS

we analyzed the wildlife diagnostic database of the diagnostic institute between 2002 and 2019 (raw data available on request). This analysis spans two decades during which diagnostic procedures have remained largely unchanged [23,24]. Since 2002, wildlife health assessments are carried out by rotating diagnostic trainees supervised by a board-certified pathologist. The largest change within the study period was the progressive implementation of health assessments in amphibians and reptiles since 2010. The grouping of the investigated taxa is shown in Table 1.

**Table 1.** Grouping of taxa applied during analysis.

| Group/taxon | Examples of frequent species | Law (mandate) |
| --- | --- | --- |
| Protected rodents | Beaver (*Castor fiber*), European squirrel (*Sciurus vulgaris*) | Hunting law |
| Hunted rodents | Alpine marmot (*Marmota marmota*) | Hunting law |
| Lagomorphs | European brown hare (*Lepus europaeus*), mountain hare (*L. timidus*) | Hunting law |
| Hunted indigenous carnivores | Red fox (*Vulpes vulpes*), Eurasian badger (*Meles meles*), martens (*Martes* spp.) | Hunting law |
| Protected indigenous carnivores | Eurasian lynx (*Lynx lynx*), grey wolf (*Canis lupus*), golden jackals (*Canis aureus*) | Hunting law |
| Non-indigenous carnivores | Raccoon (*Procyon lotor*) and raccoon dog (*Nyctereutes procyonoides*) | Hunting law |
| Ungulates | Roe deer (*Capreolus capreolus*), Alpine chamois (*Rupicapra rupicapra*), Alpine ibex (*Capra ibex*), wild boar (*Sus scrofa*) | Hunting law |
| Birds of prey | Buzzards, owls, eagles, falcons, vultures | Hunting law |
| Songbirds | Tits, blackbirds, crows, including storks, pigeons, galliformes | Hunting law |
| Waterfowl | Ducks, swans, herons, gulls | Hunting law |
| Farmed deer | Fallow deer (*Dama dama*), red deer (*Cervus elaphus*) and sika deer (*Cervus nippon*) | - |
| Domestic animals (diagnostic of predation) | Sheep (*Ovis aries*), goats (*Capra aegagrus*), calves (*Bos taurus*) | - |
| Eulipotyphla | European hedgehog (*Erinaceus europaeus*) | Nature protection law |
| Bats | *Myotis* spp., *Pipistrellus* spp. | Nature protection law |
| Small rodents | Mice (*Muridae*) | Nature protection law |
| Reptiles | *Natrix* spp. | Nature protection law |
| Amphibians | Frogs (*Rana* spp.), toads (*Bufo* spp.), salamanders (*Salamandra* spp.), newts (*Triturus* spp.) | Nature protection law |

Data analysis aimed to identify fluctuations or trends in numbers of diagnosed causes of death/diseases, submitter types and geographical origin of submitted species in the defined period. Maps depicting the origin of the cases were generated with the free software QGIS 3.20.2 (Free Software Foundation Inc., Boston, Massachusetts, USA).

#### For comparison purposes, we also analyzed the Swiss hunting statistic database between 2011 and 2019

Cantons are free in how, and how detailed, they record wildlife found sick, injured or dead, and use a variety of systems from paper trail to online reporting system. However, all cantons feed a minimum amount of information regarding carcass finds into a federal system. Carcasses found dead or with clear signs of disease are filled in the category “age, disease, weakness” (available from www.jagdstatistik.ch). Numbers pertaining to (1) hunted ungulates, (2) hunted carnivores and (3) lagomorphs (hares) were examined more closely.

### Online survey on case selection in the field

We conducted an online survey (LimeSurvey GmbH, Hamburg, version 4.3.15) with official field partners (hunting administrators, game wardens and hunters) to understand the submitters’ perspectives, needs, and practices regarding submission, health assessment, and information. We contacted the 26 Swiss cantonal hunting administrators and the corresponding person in the Principality of Liechtenstein, as well as the Swiss National Park by email and asked them to (a) fill a form themselves and to (b) forward the survey link to game wardens, park rangers or hunters that were part of their administration. The questionnaire consisted of a maximum of 46 opened and closed questions, which varied in number and content depending on given answers (S1 Appendix). The first 15 questions collected demographic details of the participants, including name, canton, district of surveillance, function, starting date of the employment in this position, professional education, attended training courses related to wildlife health, and authority for case selection. Answers were a prerequisite to proceed further. Subsequent questions focused on personal criteria of past case submissions, including numbers and incentives, with details on species and further circumstances, such as numbers of affected animals or field observation. We also asked participants to rate the current support concerning case selection (by the cantonal administration and/or the diagnostic institute), whether additional support was deemed necessary and if yes, who should provide it. The online questionnaire was open for a total of six weeks.

For descriptive analysis and statistical analysis, data were exported from the LimeSurvey software to a Microsoft Excel spreadsheet (2016 version, Redmond, Washington, USA). For descriptive statistics, median, mean, minimum and maximum were calculated for continuous variables, and percentages for categorical variables. Due to abstentions and the possibility to select multiple answers for the same question, obtained percentages did not always add up to 100%. Statistical tests were conducted using the R software [25]. Logistic regression was performed to assess associations between multiple explanatory and response variables. Explanatory variables were hunting system, distance from the diagnostic institute (distance between the offices of the cantonal hunting authorities and the center in km as indicated by www.maps.google.com), years of experience, function and level of training of the respondent. Response variables included carcass submission for carcasses with and without visible signs of disease, the frequency of annual submissions of different taxa, and the number of animals from different taxa that were found dead (1) without visible signs of disease, (2) with visible signs of a known disease or (3) with visible signs of an apparently new disease, before carcasses were submitted. Response variables were dichotomized into “yes-no” or “high-low” depending on the nature of the variable and the answers’ distribution. A complete list of all variables is available in the supporting information (S 2 Appendix). For each outcome variable, univariable logistic regression models were run for each explanatory variable. Variables with p-values <0.2 were taken as the initial variables in the multivariable model for each response variable. Further variable selection was undertaken applying a backward elimination procedure based on the Akaike Information Criterion using the “stepAIC” function of the “MASS” package [26]. The level of statistical significance was set to p = 0.05.

### Interviews with national GWHS program partners

To understand the situation and needs of national partners and stakeholders regarding wildlife health assessments, we conducted phone interviews in summer 2020. Representatives of the 26 cantonal hunting administrations and the Principality of Liechtenstein, as well as ten non-cantonal institutions (seven wildlife rehabilitation centers; two reptile and two amphibian experts from the Swiss Amphibian and Reptile Conservation Program; one representative from the Swiss National Park were interviewed. Questions were provided ahead of time (S 3 and S 4 Appendices). The aim was to assess the general satisfaction with the diagnostic institute’s services, and with the current information and communication situation regarding wildlife disease occurrence on the national and international level. Furthermore, we gathered perspectives regarding the interest in, and practical feasibility of, a potential national online reporting system for wildlife diseases. The non-cantonal institutions were additionally asked for their criteria to submit a case for postmortem investigation.

### Ethical approval

This study was reviewed by the Cantonal Ethics Committee for Human Research, Bern, Switzerland (Req-2020-00416) on 10 April 2020 with the result that it does not fall under the Human Research Act (Art. 2, 1) and therefore does not require an ethical approval. Survey participant consent was waived by the ethics committee.

### Assessment of diagnostic effort

To understand caseload-related aspects of diagnostic processes and procedures at the diagnostic institute, a time-per-task analysis was conducted. Between June 2020 and May 2021, four rotating diagnostic trainees with varying levels of experience in wildlife pathology and four supervisors documented the time required to complete certain aspects of postmortem examinations. Trainees recorded pre-/postprocessing activities, necropsy, histology, and reporting (Table 2). Supervisors reported supervision in necropsy room, reading histological slides with the trainees and report revision. The time required to perform these tasks was recorded for three types of cases (Table 3): protected species cases with in-depth health monitoring, routine cases that included histology, and routine cases without histology. Importantly, the time required for tasks not performed by trainees or supervisors, e.g., the preparation and staining of histology, the performance of ancillary test (e.g., bacteriology, PCRs) the printing and postage of reports or the archiving of samples were not recorded.

**Table 2.** Activities/duties of trainees.

| Activity | Examples |
| --- | --- |
| Preparation/ follow-up | Oral communication with submitter, opening a new case file in the diagnostic software, preparing sampling tubes, processing samples, disposal of material |
| Necropsy | Including collecting measurements and samples, taking photographs, discussion of case with supervisor, cleaning |
| Histology | Tissue trimming, slide reading, discussion of slides with supervisor |
| Reporting | Writing of report draft and corrections required by the involved supervisors |

**Table 3.** Types of cases.

| Case | Explanation |
| --- | --- |
| Protected species (e.g., lynx, wolf, etc.) with in depth-health monitoring | Postmortem examination following standardized procedure with additional tasks including radiology, photographs, morphometrics, weighing organs, systematic sampling for archiving, and histology of main organs |
| Routine cases of the GWHS with histology | Cases where the cause of death/disease cannot be defined macroscopically and/or by other ancillary tests |
| Routine cases of the GWHS without histology | Cases where the cause of death/disease can be defined macroscopically and/or by other ancillary tests |

## Results

### Case numbers

#### During the 18-year study period, wildlife diagnostic cases investigated at the national diagnostic institute almost tripled

Annual numbers fluctuated between a minimum of 132 submissions in 2002 and a maximum of 488 submissions in 2018. The current development seems stable (Fig 2A).

**Fig 2.**
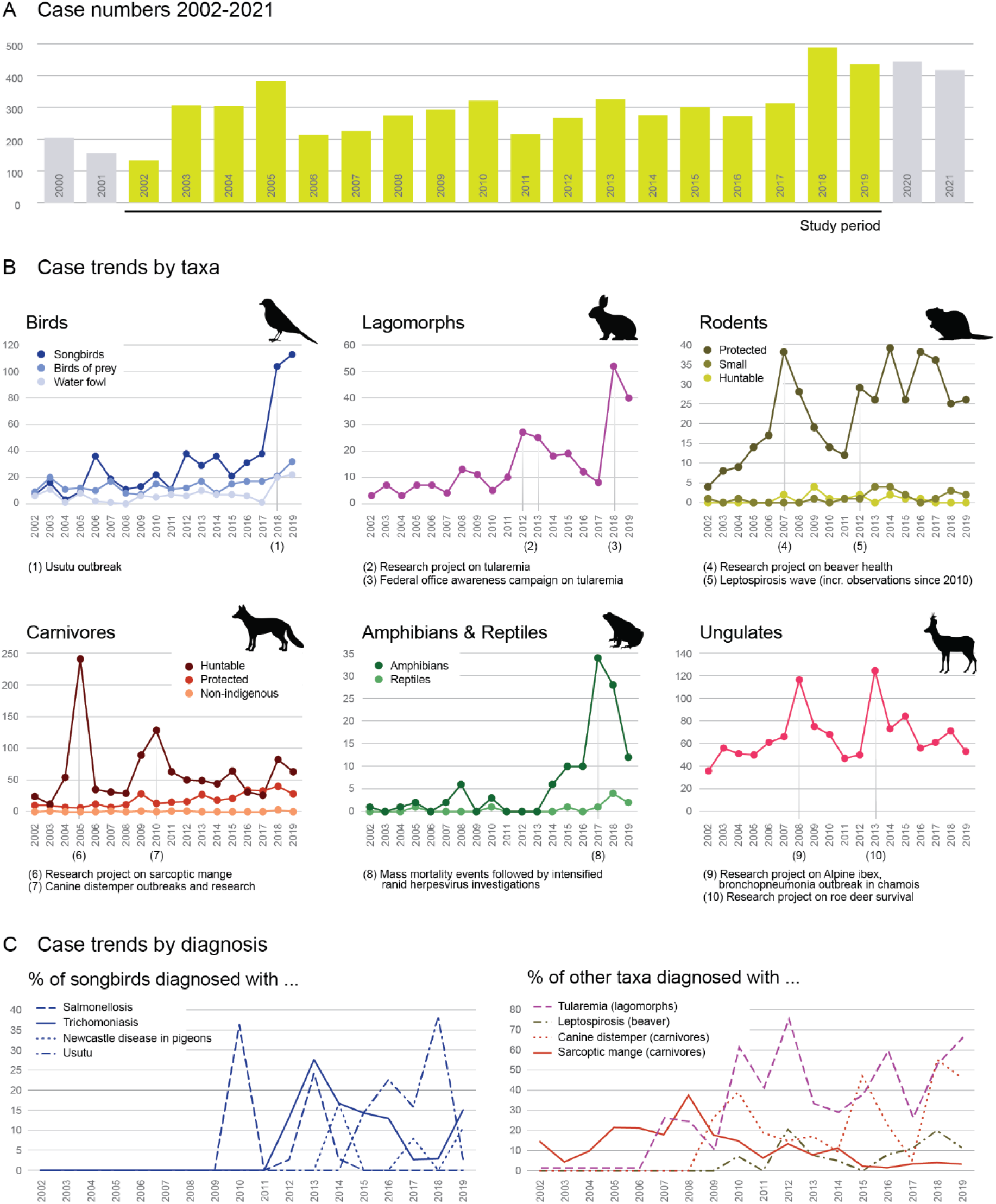
Case patterns. A) Total cases. The annual number of carcasses submitted for diagnostics rose from less than 150 in 2002 to a stable number of 450 carcasses from 2018. B) Taxon-specific patterns. Birds, lagomorphs, rodents, carnivores, amphibians and reptiles (see Table 1 for taxon grouping) feature submission increases, while ungulates remained largely stable during the study period except for two peaks in 2008 and 2013. Taxon-specific local maxima were fueled by research projects, health campaigns, disease outbreaks and mortality events. C) Pathogen-specific patterns. Similarly, specific diagnosed diseases show (mostly upwards) trends, such as Usutu, distemper, sarcoptic mange or tularemia.

#### The overall upward trend in submissions was fueled by specific taxa

Marked increases in submissions were observed for birds, lagomorphs, protected rodents, carnivores (protected and hunted) and amphibians. In 2018 and 2019, songbirds constituted about 22% (104/488) and 26% (113/437) of all cases compared to 5% (7/132) in 2002. Similarly, the number of submitted lagomorphs continuously increased since 2002 with a large peak increase in 2018. Numbers of beaver (protected rodent) submissions increased massively from 2002-2007 (2007 (17% of all cases, 38/225). After a decrease from 2008-2011, numbers fluctuated in a high state until 2017 and decreased to 6% (26/437) of all cases in 2019. Numbers of submitted carnivores (both protected and hunted) have risen constantly. Reptiles and amphibians were a relatively minor component of the submitted animal load until 2017, when an exponential increase has been observed and maintained in the subsequent years especially for amphibians. Other taxa, such as ungulates, constitute a substantial portion of submitted cases but did not contribute to the upwards trend.

#### Taxa-specific fluctuations were fueled by specific events and specific taxa network building

For birds, lagomorphs, rodents, carnivores, amphibians and ungulates, disease outbreaks, focus projects, and increased public awareness caused pronounced fluctuations. More precisely, alterations in submissions can be assigned to the following causes: birds, outbreak of Usutu; lagomorphs, targeted research project (tularemia) and media attention; protected rodents (beavers), targeted research project and first detection of leptospirosis; carnivores (foxes), targeted research projects on sarcoptic mange; amphibians and reptiles, since 2010 the submission of amphibians and reptiles was actively fostered and fueled by research projects in partnership with stakeholders (Swiss Amphibian and Reptile Conservation Program) focusing on emerging pathogens and mass mortality events investigations (conservation); ungulates, research projects (roe deer survival), bronchopneumonia outbreak in chamois (Fig 2B).

### Causes of death

#### Disease patterns and diagnosis frequencies

The cause of death in submitted birds was often undetermined (33% of all bird cases, 314/942). Four peaks of case submissions were observed due to increased disease awareness due to avian influenza in 2006, frequent salmonellosis in 2009/2010 and 2013 and Usutu in 2018 (59/104 between 2015-2019). In 2013 trichomoniasis and in 2014 Newcastle Disease of pigeons were more frequently diagnosed than in previous years (Fig 2C).

The most common cause of death of lagomorphs since 2010 was due to tularemia (109/216 between 2010-2019). In protected carnivores with in-depth health monitoring, trauma was most often observed as cause of death, while in hunted species (mostly red foxes) mainly sarcoptic mange and distemper were diagnosed. Mortality events in foxes due to distemper periodically occurred since 2009. The percentage of foxes with sarcoptic mange peaked in 2008 and showed a declining trend since then (Fig 2C). Most frequent diagnoses in amphibians included traumatic events (19/115; 17%) and infectious agents (Herpesvirus 15%; 17/115). The etiology of more than half of the investigated deaths (57%; 66/115) remained undetermined.

### Submission data

#### The diagnostic institute receives cases from diverse submitters but mostly relies on game wardens and hunters

The retrospective case analysis showed that local authorities (games wardens, hunting administrations, police, veterinary offices) constituted the largest proportion of submitters throughout the study period, with a peak of 80% in 2010 and between 60% and 70% in 2002 and 2019, respectively. Submissions from private persons, wildlife rehabilitation centers and zoos (submission of wildlife found dead/euthanized in the zoo) were subject to larger fluctuations. Contributions from private people and the Swiss Ornithological Institute have been increasing (3% to 11% and 8% to 11% between 2010 and 2019, respectively), while the contribution of wildlife rehabilitation centers declined from about 15% in 2002 to 4 and 6% in 2010 and 2019, respectively. Zoos contributed between 2 and 4% of cases throughout the study period (Fig 3A).

**Fig 3.**
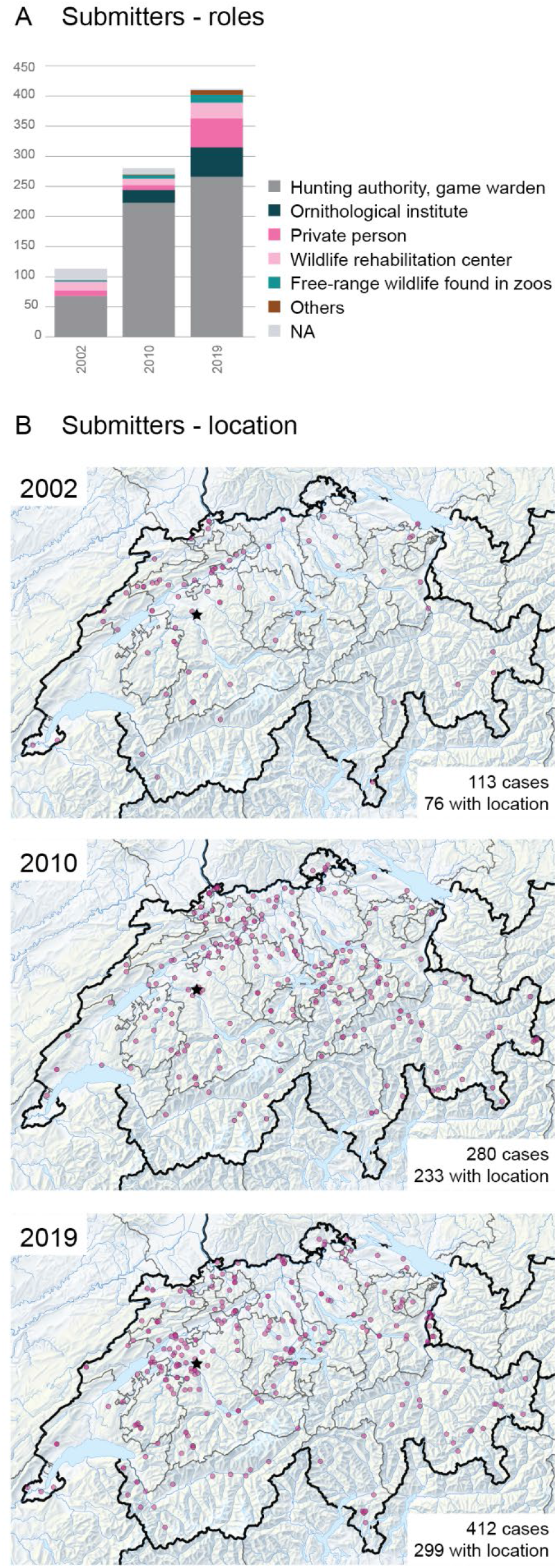
Submission patterns. A) Submitter roles. ~2/3^rd^ of all cases are submitted through hunting-associated stakeholders, while ~1/3^rd^ of cases come from other stakeholders in 2002, 2010, and 2019. The relevance of certain alternative sources is fluctuating through the years. B) Submitter location. Between 2002 and 2019, the geographic source range of the submissions expanded to previously uncovered regions and administrative units.

#### Biogeographical and political boundaries matter for case submissions

The geographic origin of the submissions was mostly concentrated on the northwestern part of the country in 2002, but the spatial case distribution spread to the eastern and southern parts of Switzerland in 2010 and 2019 (Fig 3B). Statistical analyses of the online survey revealed that the distance to the diagnostic institute was not related to the submissions of carcasses, neither in general nor for those with or without visible disease signs. The distance from the diagnostic institute had no effect on the number of animals of any taxa submitted per year, except protected rodents, for which numbers decreased with the distance (p = 0.04, OR = 1.00, CI = 1.00-1.00). However, data from the retrospective case analysis suggest that distance still plays a role as carcass submissions from distant cantons are sparse (Fig 3B).

#### Only a fraction of discovered carcasses or diseased animals are submitted to the GWHS

On average, between one and three percent of animals reported in the Swiss Federal Hunting Statistics in the category “age, disease, weakness” were submitted to the GWHS between 2011 and 2019 (Fig 4A). The proportion of submission is species dependent. For example, between 15% to 75% of lagomorphs found dead were examined at the diagnostic institute with a marked increase since 2018 (Fig 4 A). The retrospective case analysis demonstrated that not all taxa are represented equally in all geographic areas. Data from 2002, 2010 and 2019 together illustrate that ungulates and carnivores (hunted and protected species) are submitted from all over the country, while lagomorphs, rodents, eulipotyphla, amphibians and reptiles (not hunted) are mostly submitted by areas north of the alps. (Fig 4 B).

**Fig 4.**
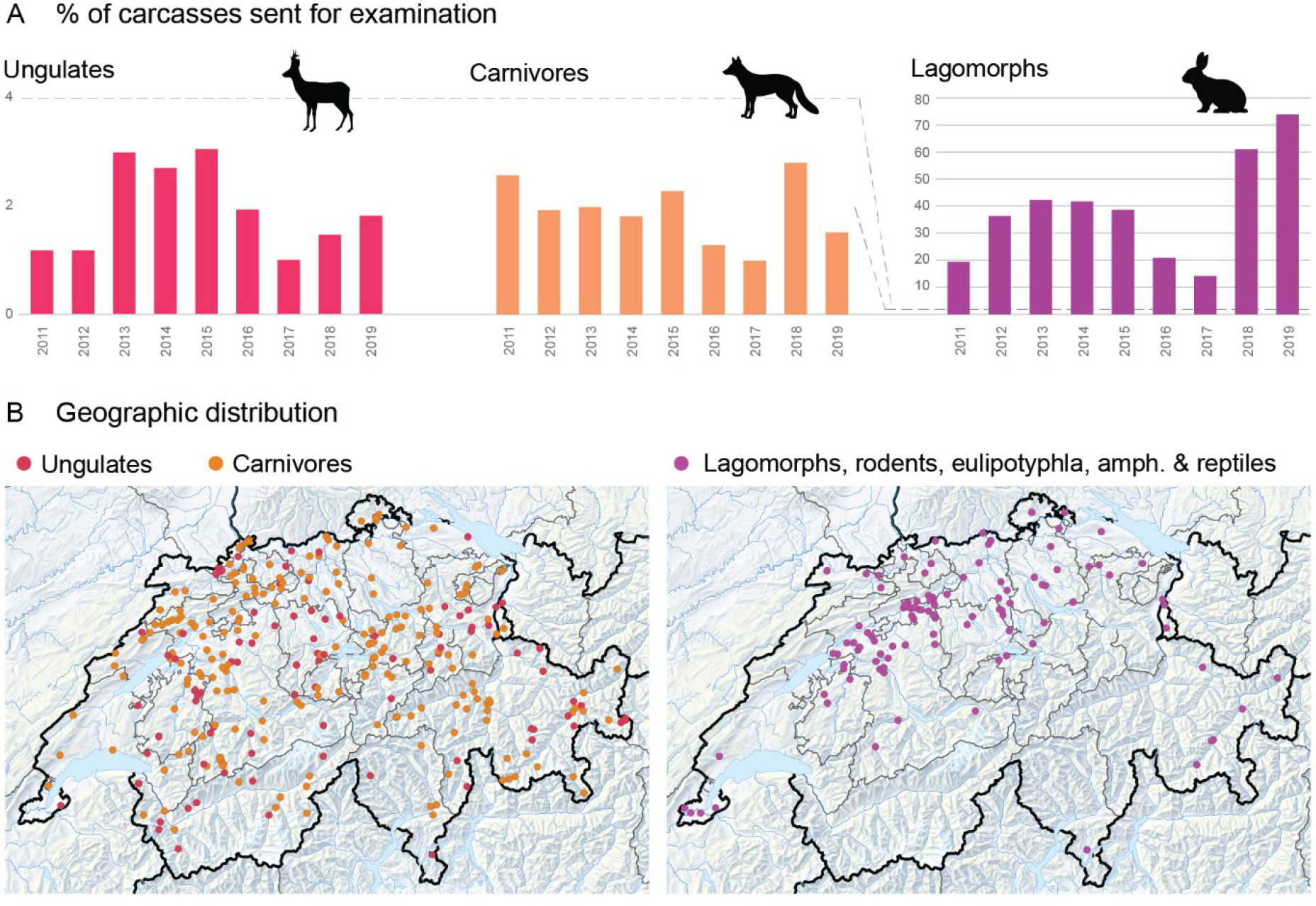
Submission gaps. A) Taxon gaps. Only ~3% of ungulates and carnivore carcasses are submitted to the GWHS for examination. For other groups, such as lagomorphs, the numbers are much higher thanks to a) federal information campaigns and b) research-driven requests to submit lagomorphs for tularemia testing. B) Geographic gaps. Not all taxa are represented equally in all geographic areas. Data from 2002, 2010 and 2019 together illustrate that ungulates and carnivores (hunted and protected species) are submitted from all over the country, while small species (not hunted) are mostly submitted by areas north of the alps.

#### The online survey had a high response rate with most of the responders being professional game wardens

We received a total of 238/356 complete questionnaires from 25/26 Swiss cantons as well as the Principality of Liechtenstein, which corresponds to a return rate of 67%. Forty-two percent of the responses were from cantons with a hunting ground system (100/238), 54% from cantons with license system (130/238) and 3% from the canton of Geneva, where hunting is prohibited (8/238). Most participants were professional game wardens (54%, 129/238). The respondents had been in their current position between one month and 42 years (median: 10 years). Most responders had a trade profession as primary education (43%, 102/238). Thirty-eight percent had completed a federal game warden training (90/238) in addition to the cantonal formation. Many others (69%) had taken courses regarding wildlife health (165/238), game meat hygiene (65%, 155/238), wildlife diseases (53%, 127/238). Few had taken courses regarding diagnostic of predation (2%; 4/238) and wildlife immobilization (4%; 1/238) or other courses (6%; 15/238) and 6% have neither completed any courses nor a federal game warden training (15/238).

#### Cases are submitted to understand the cause of death

Participants indicated they would submit carcasses upon suspicion of an infectious disease (85%; 202/238), to assess the cause of death/disease (62%; 149/238), or upon suspicion of predation (32%; 75/238). Furthermore, half of all responders (55%; 131/238) have already sent dead animals with unknown cause of death and no visible signs of trauma or disease. Carcasses with lesions or clinical signs typical of a specific disease, e.g., sarcoptic mange or distemper, were submitted by only half of all respondents (56%; 134/238), mostly to confirm the field diagnosis (92%; 123/134).

#### Cases are also submitted to satisfy monitoring needs and directives

The online survey participants indicated they would submit carcasses to rule out poaching (22%; 53/238) or hunting outside hunting season (27%; 64/238). A total of 168/238 participants (71%) from 21 cantons indicated that some species were monitored for specific diseases in the framework of targeted health surveillance programs in their canton, and this was also a major reason why carcasses with typical disease signs (see above) were nonetheless submitted to the diagnostic institute or to specialized monitoring programs elsewhere (41%; 55/134). Targeted species and diseases listed by the respondents included wild boar (trichinellosis, African/classical swine fever), brown hares (tularemia, myxomatosis), red deer (tuberculosis, foot rot), waterfowl (avian influenza, Usutu and Newcastle disease), beaver (leptospirosis), fox, martens and badgers (distemper). In line with this, participants most commonly submitted hunted carnivores (63%; 30/48), followed by ruminants (44%; 21/48), protected carnivores (35%; 17/48) and lagomorphs (29%; 14/48).

#### Further reasons for submissions

were specific interests of the submitter, e.g., the exclusion of a certain infectious agent (26%; 35/134), and specific requests of the diagnostic institute hosting the GWHS to receive carcasses with suspicion of a specific disease, e.g., for teaching purposes or research projects (18%; 24/134).

#### Cases were not submitted when the infectious cause of death was obvious, or if death was caused by trauma

Responders did not submit carcasses with visible signs of disease if a) they trusted that their field diagnosis was correct (65%; 66/102), e.g., sarcoptic mange, distemper, infectious keratoconjunctivitis; if b) they had received instruction from their authorities, e.g., fox with suspicion of mange, (13%; 13/102) or c) if there were no more requests from laboratories/institutions (3%; 3/102). Trauma cases are also usually not submitted in species with no in-depth health monitoring. A total of 174 out of 238 participants (73%) stated that they have not submitted any carcass with clear evidence of a trauma (e.g., traffic accident) so far. Seven percent (16/238) answered that they send 1-2 animals with signs of trauma per year and 7% (16/238) said they send 5-10 such animals per year. Five percent (12/238) stated 3-4 animals per year and 2% (4/238) send more than ten per year.

#### Case selection for submission is related to education and experience

The experience as well as completion of continuous education courses and the primary education were detected as factors influencing the case submission. Game wardens who completed continuous education courses and also had knowledge from their former profession (e.g., butcher, farmer, forester) (S 2 Appendix) stated significantly more often that they had already submitted animals for postmortem investigations compared to responders who only had experience through their former profession (p < 0.001, OR = 0.73, CI = 0.62-0.86) or game wardens without additional training (p = 0.01, OR = 0.29, CI = 0.14-0.62). Most respondents (87%; 207/238) were authorized to decide on their own whether a carcass shall be sent for postmortem examination. If not authorized, decision depended on the hunting authorities (81%; 25/31).

#### Case selection in the field encompasses a limited extent of epidemiological considerations

According to the case records, many submissions are declared as a single animal problem as opposed to a population issue. Sensitivity for epidemiological issues depends on the species, and on visible signs or indications for infectious disease. The majority of the responders stated that in case of increased mortality within the same month and district without suspicion of a specific cause of death they would immediately submit a carcass for investigation only if the concerned species was wild boar (36%; 87/238), raccoon or raccoon dog (29%; 70/238) or a protected carnivore (65%; 157/238). For other taxa, 18% (70/238) answered that a minimum of 2-3 carcasses was required prior to first submission. If increased numbers of dead animals with lesions or signs indicative of a suspected disease which is already known within the area were observed, the same answers were given as for animals without signs of disease. Only if protected rodents (beaver) were involved most participants (23%; 54/238) would send the first carcass found for examination. For the same scenario, except with the suspected disease being new to the area, the majority of responders would send the first observed carcass for further investigation, except for hunted carnivores, waterfowl and eulipotyphla. In those cases most responders, meaning 31% (73/238), 23% (55/238) and 19% (46/238) would wait until 2-3 dead animals were found within a month, respectively. For ruminants, submitters waited significantly longer to submit (i.e., until more dead animals without visible signs of disease in the same area within a month occurred) the more distantly they were located from the diagnostic institute (p = 0.007).

#### Hunting systems also affected case selection

For example, wild boar were sent significantly more often from the canton of Geneva (state hunting system) than from cantons with license system (p = 0.02, OR = 1.67, CI = 1.21-2.29). Regarding protected carnivores, respondents from cantons with hunting ground system sent significantly less animals per year than those from cantons with license system (p = 0.04, OR = 0.85, CI = 0.72-0.99).

#### The availability of local veterinary expertise also impacts submission numbers

The majority of respondents who had already submitted carcasses to an institution/laboratory for postmortem examination (77%; 184/238) submitted to the diagnostic institute (70%; 167/238). The other cases were sent to other cantonal or academic laboratories. If responders had not submitted any carcass for necropsy yet, reasons were mostly because they had not had any case of interest (77%; 36/47), they could discuss cases with their local veterinarian (15%; 7/47) or the next laboratory/institute was too far away (4%; 2/47). Almost half of responders indicated that they did contact the cantonal veterinary office before carcass submission (47%; 111/238). Consultation of the cantonal veterinary office prior to carcass submission was done only in case of a suspected reportable disease by 62% (69/111), while 25% (28/111) said that they always do it. Of the responders who do not contact the veterinary office prior to carcass submission, 9% (11/122) contact the office after case submission and 38% (45/122) report to the veterinary office only if a notifiable disease is diagnosed.

#### Subjectively perceived increases or decreases in submission numbers were attributed to a variety of reasons

The majority of responders (77%; 184/238) answered they had submitted similar numbers of animals annually in their current function, while 8% (20/238) thought that the number had increased and 8% (19/238) that it had decreased. Six percent (15/238) did not answer the question. The increase was justified in 65% (13/20) as secondary to higher awareness for diseases (e.g., African swine fever, tularemia, avian influenza and tuberculosis), 55% (11/20) to growing populations of protected species (e.g., lynx, beaver, wolf) and 45% (9/20) to the perceived increase of dead or diseased animals (e.g., beaver and orphaned lynx). Main reasons for decrease included a decline of the observed morbidity and mortality (53%; 10/19), (e.g., sarcoptic mange, distemper, rabies), or changed hunting administration instructions (42%; 8/19). Furthermore, the longer a responder had been in service, the more likely they had the impression of decreasing submission numbers (p < 0.001), while there was no relation between the years in service of the responder and the rating of case submissions as increasing (p = 0.56). The complete list of statistical data can be found in the supporting information (S 5 Appendix and S 6 Appendix).

#### Interviews and online survey clearly indicate that a successful implementation of a nation-wide online reporting system requires addressing several concerns

Phone interviews with the cantonal authorities revealed that fourteen cantons (52%) already work with an online reporting system (ORS) that allows registering wildlife disease occurrence (at various levels of detail). Five additional cantons are planning to set up a digital/app-based system for wildlife management which includes disease recording in the near future. The remaining eight cantons do not have any kind of ORS or do not record wildlife diseases at all. In line with this trend, the majority of hunting authorities (85%; 23/27) and non-cantonal institutions interviewed (100%; 10/10) considered the option to register animals found dead/diseased, but not submitted for pathological investigation useful. Many indicated that such a system would give a better overview of the occurrence and spread of wildlife diseases in Switzerland, and that it could be a useful tool also in connection with existing hunting statistics. While 53% (126/238) of respondents of the online survey did not see an added benefit in a nation-wide ORS, 43% (103/238) expressed views in favor of such a system. Concerns included (1) fear of redundancy and increased efforts. Several electronic systems to record dead wildlife are already in place, and stakeholders were wary of redundant efforts to record cases into a second system. A system that increases administrative effort without yielding substantial benefit at the local level was seen as problematic. Also, concerns focus on (2) reliability. Respondents were concerned that the records would be of questionable informative value as the diseases would only be diagnosed in the field by non-veterinarians, based on external gross lesions, i.e., bearing a risk of misdiagnoses due to the omission of a pathological examination by an expert. Finally, concerns related to (3) the technological requirements of an ORS. Given that ORS require the ability to use an app on a smartphone, respondents pointed towards these requirements as a potential challenge for older hunters.

### Stakeholder’s needs

#### The WHS operates in a fragmented legislative space

Various national laws warrant the monitoring of wildlife diseases, and international agreements oblige to monitor the epidemiological situation of wild animals. The GWHS is based on legislation stating that the federal government should support wildlife disease investigations [27]. First, the animal disease law defines animal diseases to be controlled or eradicated that also affect wild animals and regulates the role of game wardens and hunters and the involvement of official veterinarians. Second, at the level of species protection legislation, the federal government has a responsibility to promote research on wild animals, their diseases and their habitat through the hunting law. For wildlife species covered by the nature protection law, the mandate for disease monitoring is not as clearly formulated. In circumstances that are legally not entirely clear, personal interest, passion and initiative play a significant role. An excellent example for this are amphibians and reptiles. These were originally not covered by the diagnostic institute (given its original focus on hunted species), but personal interest of a new hire led to increasing collaborations and case numbers, which led to federally financed research projects, and finally (in 2020) to an official mandate for amphibian and reptile health monitoring.

#### The GWHS operates in a field of partly contrasting needs of the main funding bodies

The FOEN focuses on population level aspects and insists that general wildlife health surveillance offered at the diagnostic institute meets the legal requirements and the needs of the involved partners concerning pathological investigations, information on current disease dynamics, and education. Largely, the focus of the FOEN relies on species protected by the hunting law. The hunting administrations focus on wildlife management, expect advice in management actions and request case-based support. The FSVO and cantonal veterinary services, in contrast, focus on early warning for emerging diseases, reportable animal diseases and zoonoses, and are thus interested in individual-level data. They do not have a species focus. Finally, the University of Bern (where the diagnostic institute is integrated) focuses on research and teaching tasks. Accordingly, the diagnostic institute needs to maintain, in parallel, a high quality of wildlife health surveillance and diagnostics, teaching activities (requirement by the university for veterinary student education and by the authorities for field partner training), and continuous education of the diagnostic institute’s staff (expertise maintenance).

#### The diagnostic needs of local administration and submitters are currently mostly met

Overall, hunting authorities in Switzerland and Liechtenstein as well as the non-cantonal institutions expressed satisfaction as concerns the services offered by the diagnostic institute concerning wildlife health surveillance. They submit the majority of wildlife carcasses where they a) want to know the cause of death or b) have the suspicion of an infectious disease but cannot determine the disease or c) suspect intoxication. A service which could be expanded, according to stakeholders, is toxicological investigations (i.e., specific laboratory analysis in cases of suspected poisonings), and amphibian health surveillance.

#### Support needs of submitters are also mostly met, and open needs mostly concern cantonal authorities

According to the online survey support from cantonal authorities for decision on case submission was rated as “very good” (55%; 132/238), “good” (28%; 66/238) or “sufficient” (11%; 27/238). Only 1% (3/238) answered “bad” and only one respondent rated it as “very bad” (1/238). The majority (76%; 182/238) stated that they did not need any additional support in this matter from any institution. Thirteen per cent (30/238) named the cantonal veterinary authorities as institution they would like to have more support from, 12% (28/238) from the cantonal hunting administration, 8% (20/238) from the diagnostic institute, 4% (10/238) from the FOEN, 2% (4/238) mentioned the FSVO. One person each mentioned a private veterinarian and the cantonal laboratory.

#### The GWHS does not satisfy non-cantonal stakeholder’s information needs

While the needs of cantonal authorities regarding wildlife disease investigation and information were covered and also benefited from regular inter-cantonal and even international interactions, non-cantonal institutions were less satisfied with the availability of information on cantonal and national level. Information requirements were related to biogeography (desire for an online platform or map) and to information access on the local, federal and international level. For example, 8 of 10 non-cantonal institutions stated that they did not have sufficient information about the cantonal and national wildlife health situation, and four did not have enough information about disease developments and epidemiological trends in Europe.

### Assessment of diagnostic effort

#### Time-tracking of various tasks associated with diagnostics uncovered three factors contributing to time per case

**Firstly**, and most importantly, detailed health monitoring requirements for protected species was time-consuming. Postmortem analyses of protected species took about six hours, with a twice-as-long necropsy compared to routine cases. The required amount of time for supervisors was approximately 90 minutes for protected species compared to 60 minutes for routine cases. **Second**, histological examinations extended the time per case. Routine cases with histology took about three hours compared to one hour for cases without histology. Histology also added 30 minutes of supervisor time. **Finally**, the academic teaching setting with the associated high personnel turnover and the high fraction of trainees contributes to time per case since variations in the data correlated with trainee experience. However, given the lower cost of trainees, longer time per case due to the academic setting does not translate to higher resource needs. Also, tasks that do not require veterinary training (e.g., case entry into the system, specimen processing, necropsy preparation, cleaning of necropsy hall) took approximately 17% of the time a veterinarian needed to complete a case (23/139 min).

### Current gaps of the WHS

#### Our analyses revealed gaps in the GWHS regarding specific species and particular diseases

**Firstly**, common or easily recognizable diseases are underrepresented in the monitoring dataset. For example, sarcoptic mange is usually recognized by practitioners in the field, and therefore not submitted. Between 2002 and 2019, a total of 179 cases of sarcoptic mange were recorded, with only 10 submissions from 2015 onwards (Fig 2C), which clearly underrepresents the true prevalence of the disease among the Swiss carnivore population. **Secondly**, the focus of the GWHS on the hunting and game system causes blind spots in species that are not covered by the hunting law, that are not relevant in a hunting context, or that are only occasionally hunted. For example, reptiles, amphibians, bats, eulipotyphla, small rodents and raccoons/raccoon dogs were not named among commonly submitted animal groups by the survey respondents. **Thirdly**, biogeographic aspects and habitat accessibility play a role in submissions. While detection of fresh wildlife carcasses is an intrinsic challenge of any GWHS [14,28], this is particularly challenging for remote alpine areas where access as well as submission logistics, e.g., cooling and shipment, can be difficult. Therefore, alpine species such as marmots (*Marmota marmota*) or alpine snow grouses (*Lagopus muta*) are almost non-existent in the GWHS, even though they are not rare.

#### In addition to the gaps in the data, our analyses reveal three types of issues with regard to data access

**Firstly**, a large amount of data on wildlife health is collected locally. Reporting systems range from paper notations through Microsoft Office lists to app-based reporting systems, and the ability of systems to interface with other animal and health databases varies from zero to full integration. There is no central collection or consolidation of this wealth of information. **Secondly**, specialist disease centers responsible for targeted surveillance schemes currently report to the FSVO in real-time, but this information is only shared by monthly update for reportable diseases and with even bigger delay for other diseases such as tuberculosis or trichinellosis, and only within administration and not between diagnostic centers. This leads to situations where several centers independently start to suspect a disease outbreak. **Thirdly**, currently no wildlife health data is shared systematically between the WHS (targeted or general) and species focus centers, such as the Swiss Ornithological Institute or the Bat Foundation Switzerland. The exchange of health data between centers, a prerequisite for an integrated wildlife monitoring [11], currently relies on personal acquaintance and communication, with the exception of amphibians and reptiless (official case reporting and annual summary of diagnoses since 2017).

## Discussion

This study evaluated submission practices and case trends in the framework of a European GWHS, with the aim to better understand (1) the impact of disease trends, (2) current case selection practices, (3) overlapping and contrasting needs of stakeholders, and (4) factors impacting the ability of the GWHS to function efficiently and effectively. It uncovered gaps and develops recommendations on how to set up or improve wildlife health surveillance programs that face real life challenges and settings.

### Methods

#### The present study used a multi-tiered approach to understand the current characteristics, strengths and gaps of a small-country European GWHS

To our knowledge, this is one of the most exhaustive evaluations of a GWHS performed to date. We combined semi-quantitative structured interviews (local administration), an online survey (field experts), a 20-year case analysis, time tracking, and a consultation of the legal frameworks. Response rates in interviews and surveys were excellent, and together with the clear trends in the answers, and the consistent results between independent approaches, the conclusions drawn can be considered valid.

### Case numbers and causes of death

#### In summary, the data suggest that a combination of - individually minor - changes can push a GWHS to the borders of its capacities and thus its ability to perform its true role in early detection and surveillance

We found that the observed increased caseload could not be attributed to any singular cause, but was associated with infectious disease dynamics, increased public awareness for specific diseases, research activities and increasing population size of protected species.

#### Disease related factors included the emergence of novel pathogens as well as increased prevalence of known diseases

Songbirds have been increasingly submitted likely in relation to increased disease awareness due to an avian influenza epidemic in 2006 as well as mortality events associated with infectious diseases such as salmonellosis (2010 – 2013) [29] or the emergence of Usutu (2018) [30] in Europe. In hunted carnivores, case increase was mainly due to infectious disease spread, such as sarcoptic mange [31] and distemper [32,33]. Distemper simultaneously emerged in surrounding countries and spread up to Denmark [34].

#### Societal factors were related to public perception as well as education

The most submitted species, such as roe deer, red deer, chamois and ibex, are appreciated game species and benefit from increased attention by hunters and hunting authorities. Also, roe deer and foxes are widely distributed throughout the country [35], and often appear close to human settlements. This increases the likelihood of carcasses to be found, which is a typical challenge in wildlife health surveillance [14,28], but also bears a risk of pathogen transmission to livestock, domestic animals or even humans [36–38]. Generally, however, field experts base their case selection on valid criteria, such as the species, suspected cause of death (infectious vs non-infectious) and apparent local disease emergence. Additional training like the completion of continuous education courses resulted in a higher wildlife disease awareness, underlining the positive effect of training field partners on wildlife diseases. Total number of submitters increased which is illustrated by the higher proportion of cantons from which cases originated.

#### Research projects on focus species, inspired by perceived changes in prevalence or by stakeholder’s needs, sometimes had long-term impacts on submission practices

Research projects contributed to increased submissions of **lagomorphs** (especially European brown hares) for a study on tularemia (2012-13) [39]. Compared to a former study on hare diseases [40], tularemia occurrence significantly increased within 13 years and high case numbers have continued to be recorded since then. This was shown by the high percentages of lagomorph submissions compared to the total number of diseased animals recorded in the Swiss hunting statistics and the marked increase of hare submissions since 2018. High numbers of mangy **foxes** were related to a research project in 2003-05 and to targeted surveillance activities over nearly the whole study period [31,41]. The increase in **amphibian** submissions since 2017 was due to the setting up of a herpetology network and amphibian health surveillance program, including investigations of mass mortality events [42,43]. Another reason for increased case submissions were new partnerships, for example with the Swiss Ornithological Institute since salmonellosis detection in passerine birds in 2010, and with the Swiss Amphibian and Reptile Conservation Program since 2013.

#### Finally, increasing population sizes can present unexpected challenges to a GWHS

Increased case submissions of **protected species** were likely most often driven by population growth of Eurasian lynx, grey wolf and beaver [44–46]. Since for protected carnivores, such as the lynx, wolf and brown bear, Swiss management plans require a full postmortem investigation including non-health related indicators, this increase burdens the system while not contributing to the GWHS. Similarly, submission of wild cats is strongly encouraged [8]. Beavers had an intermediary status, with the same procedure as for wild cats, but enhanced because of a research project in 2006 [47,48]. Also, **population regulation** can impact the GWHS. For example, wild boar were sent significantly more often from the canton with state hunting than from cantons with hunting permit system, which is likely due to the fact that the canton of Geneva used to have one of the highest wild boar densities [49] and local game wardens carry out population regulation.

#### Together, these results indicate that many small factors can collectively challenge a GWHS, slowly, over time

Accordingly, changes in disease dynamics, in education and awareness, in focus species, and in population sizes need to be accounted for when projecting the capacities and resources of a GWHS. This is particularly relevant if resources are dependent on longer-term cycles (in the presented case, a thorough re-evaluation is possible every four years). It is also relevant to consider development processes, such as slow geographic expansion. For example, the wildlife health surveillance in the form practiced today has been in the process of development over the last 20 years and may not yet represent a steady state system. Therefore, a combination of early-on definition of clear aims regarding the mandates of various stakeholders is key, as are regular discussions regarding implementation and execution of these mandates in view of current situations and developments.

### Submission data

#### For GWHS with a strong focus on hunted species, the value of submissions from private persons, wildlife rehabilitation centers and species centers are sometimes questioned

However, so-called garden wildlife (species such as hedgehogs or songbirds) is almost exclusively submitted by these submitter groups [50], and also not recorded in the national hunting statistics database. Diseases such as Usutu, or salmonellosis, are frequent in garden wildlife, which is usually in close contact with pets, farm animals, and humans. Promoting the submission of these non-hunt-related groups to the GWHS is key to early disease detection and monitoring in the context of a One Health perspective.

#### In a multi-stakeholder scenario, communicable, clear criteria might facilitate case selection

Importantly, the cases selected for submission and accepted for workup need to satisfy the needs of the GWHS, of specialist surveillance programs, of teaching and education, and species protection programs, and need to allow for the “known unknowns” or unexpected development. Therefore, we suggest the development of careful case definitions and a decision tree that is tailored to the goals of a general surveillance program for wildlife health, to the expectations of the mandating national authorities and the University, as well as the needs of field partners. An example developed and used at the diagnostic institute is provided with the supporting information (S 7 Appendix).

#### The GWHS features pronounced information gaps which are region- and species specific

Less than 3% of deaths in hunted and protected species are presently submitted. Importantly, the surveillance gap concerns a) rare or poorly detectable species and b) diseases that are common or easily detected. The former can be covered by targeted research projects, or “focus species” efforts. The latter gap could be covered by improved information collection and management. Again, similar to garden wildlife, a diverse group of submitters and involved stakeholders is particularly important with regard to species and disease gaps. Fostering the development of such groups must take socio-cultural and geographic aspects into account. While the geographic origin of cases gradually extended throughout the country in the study period, the three major southern cantons are separated from the diagnostic institute by the alpine crest, and deliveries of carcasses by car requires time and serious effort.

#### Another question when improving or building GWHS is whether to rely on a single diagnostic center or multiple dispersed centers

The majority of wildlife carcasses were submitted to the diagnostic institute despite the presence of other institutions that perform veterinary pathological examinations. The majority of responders indicated that the veterinary authority was not informed before case submission and many of them did not contact the veterinary authorities at all. At the same time, the Swiss Ordinance on Epizootic Diseases requires since 2017 that hunters and game wardens inform an official veterinarian in case of suspicion of a notifiable disease in a free-ranging wild animal [51]. The role of local veterinarians is therefore mixed, and likely still evolving. On the one hand, cantonal veterinary institutions primarily work with domestic animals and often lack expertise in wildlife health. On the other hand, including local veterinary expertise more in case selection, or in a notification system for systematic data collection, might be advisable to close some gaps of the current GWHS.

### Stakeholder needs

#### Serving many masters is both a challenge and an opportunity for a GWHS

The analyzed system is located at the interface of distinct funding bodies and distinct areas of responsibilities. Somewhat counterintuitively yet typical for multi-stakeholder settings in public good areas, resources to fulfill its role need to be predicted, budgeted, and negotiated by the GWHS itself. The presence of different funding sources leaves the GWHS in the position to negotiate and balance the stakeholder’s needs and ensure that everyone’s needs are met without compromising each other’s resources. Also, external funding comes with reporting obligations that are resource intense. At the same time, the situation is a win for each stakeholder. None of them could satisfy their legally mandated needs on their own without substantially increasing invested resources, or even potentially competing, e.g., cases for teaching vs diagnostics. A make-or-break point of such a system is continuous exchange and trust among the partners to avoid personnel turnover, loss of identification with the greater cause, quiet quitting, or retraction from one partner from the construct. Another make-or-break point is access to highly motivated broadly skilled individuals that can teach, diagnose, report, account, and strategically plan while being at the forefront of wildlife health research. This is particularly challenging in the context of a university pay grade system that a) does not offer attractive wages and b) offers limited long-term career options for mid-career scientists.

#### While most needs are currently met by the GWHS, systematic, timely and centrally accessible information on the occurrence and prevalence of wildlife diseases is lacking

While clear processes and central information systems exist for livestock diseases, there is no central information record in Switzerland for the occurrence of non-notifiable and common diseases (e.g., distemper, sarcoptic mange). As a result, the information situation on the occurrence of those diseases is perceived as poor, especially by stakeholders of animals rarely examined at the diagnostic institute. Solutions to this information gap are discussed below.

### Demanding objectives

The combination of multiple, sometimes diverging needs also leads to a situation where certain aspects are both a blessing and a burden. Both aspects presented below may impact the diagnostic efficiency. From a scientific perspective, however, they are a treasure trove of otherwise unavailable information. For example, the processing of trauma cases of protected species requires a careful and continuous balancing of extracting the maximum of information when case numbers are low, and reducing efforts when case numbers rise. Here, the ability to immediately react and take decisions together with all stakeholders is key for the functioning of a diagnostic institute.

#### In-depth monitored protected species

(wolf, lynx, wildcat, golden jackal, beavers) present a particular challenge for the diagnostic institute. They are time-consuming and take almost three times as long as routine cases. Reasons for this are that postmortem investigations of these species include x-rays, the collection of morphometric data of the carcass and the inner organs, as well as organ sampling for histopathology and biobanking. Often, they also have to be dissected in a carcass-preserving manner for voucher and museum preservation. The academic teaching setting with the associated high personnel turnover and the high fraction of trainees contributes to time per case since variations in the data correlated with trainee experience. However, given the lower cost of trainees, longer time per case due to the academic setting does not translate to higher resource needs. Minor reductions on time per case are possible through efficiency-enhancing measures. However, this does not address the main issue for the GWHS: a large proportion of these carcasses do not contribute to the focus of the GWHS, as they often represent healthy individuals killed by trauma. Nevertheless, the apparent prevalence of pathogens found in these species is much closer to the real prevalence compared to any other taxa examined. Any reductions in efforts are annihilated by the increasing population sizes of protected species in the area. We therefore recommend that for any species with in-depth monitoring plans for increases in abundance, and upfront decisions on case numbers exist. This is essential to avoid overwhelming a system designed to monitor diseases with healthy individuals, potentially by separating capacities for health monitoring and protected species monitoring.

#### Histopathology

also contributes substantially to time spent per case. However, it has a number of advantages crucial to disease surveillance, including the detailed classification of lesions, the detection of tissue alterations that are not visible macroscopically [52] and the demonstration of associations between pathogen identification and tissue lesions [24,42]. In addition, in the academic setting of the described GWHS, histopathology – decision, sample preparation, and slide evaluation – is a key part of personnel training, in particular of post-graduate students. Carefully weighing the benefits of histopathological examinations in relation to the relevance of the case for wildlife health surveillance (individual animal vs. population problem), questions of the submitter (animal welfare, zoonosis), informative value of the additional information obtained (fresh condition of the carcass) and added value for teaching is therefore warranted to account for the priorities of all stakeholders and funders.

### Gaps

In line with previous studies [23,24], there are large, and species-specific, discrepancies between animals found dead or diseased according to the Swiss hunting statistic database and the number of cases investigated in the context of the Swiss GWHS. The most likely explanation is that diseases with overt signs such as infectious keratoconjunctivitis in wild ibex and chamois [53], sarcoptic mange in carnivores [31] or foot rot in ungulates [54], and other obvious causes of mortality (e.g., traffic accidents) are typically diagnosed in the field. Submitters specified that cases with obvious cause of death or if death was caused by trauma, were not submitted which supports this statement. However, field diagnoses yield the risk of missing underlying conditions, e.g., a fox with trauma as final cause of death might have been weakened by an infectious disease, e.g., distemper, which made him prone to be hit by a car. Cantons are at liberty to record these cases as they see fit and report them to the national hunting statistics database as “age, disease, weakness” or “traffic”, without further indication of disease etiology. This leads to the situation that wildlife health experts face a lack of accessible comprehensive overview of wildlife disease occurrence on the national level, even though the data is actually recorded to some extent at the local level and causes an information gap within the GWHS with regard to certain, often frequent, diseases. This is even more true for species not included in the legal scope of the cantonal hunting administration, where no collection of data for the national hunting statistic is obligatory. Even though the data is often collected by species-specific institutions (e.g., Swiss Ornithological Institute), it currently does not feed into the WHS.

### Measures

#### Practice-science collaborations, such as represented by GWHS programs, are capable of flexible incorporation of change

Fig 5 illustrates how different types of information contributors have distinct types of expertise regarding wildlife health. Integrating method-driven, theory-based, and validated scientific knowledge with real-world, action-based, and contextualized experimental knowledge is a key aspect of sustainable system transitions and system changes [55]. We recommend that people in charge of accompanying such change processes are familiar with basic concepts of transdisciplinarity [56] its application in multidisciplinary and multi-stakeholder policy contexts [57] and factors that increase the success chance of such projects [58]. Incorporating learnings from other fields that navigate the knowledge-to-policy line and that are forced to continuously incorporate past, present, and predicted change such as maritime spatial planning [59] or groundwater policy [60] may be useful. For example, successful policy change processes in the sustainability context tend to (i) engage stakeholders in a participatory process that includes collaborative modeling and social learning; (ii) provide improved understanding of evolving scenarios from all perspectives relevant to the topic; (iii) acknowledge and address uncertainty in scientific knowledge as well as the diversity of stakeholder preferences, e.g. using multi-model uncertainty analysis; (iv) adopt transdisciplinary monitoring and evaluation methods from the very start [60]. Other factors that play a role are (1) the maturity of relationships within the collaboration, (2) the level of context knowledge present within the collaborative team, and (3) the intensity of the engagement efforts within the project [58].

**Fig 5.**
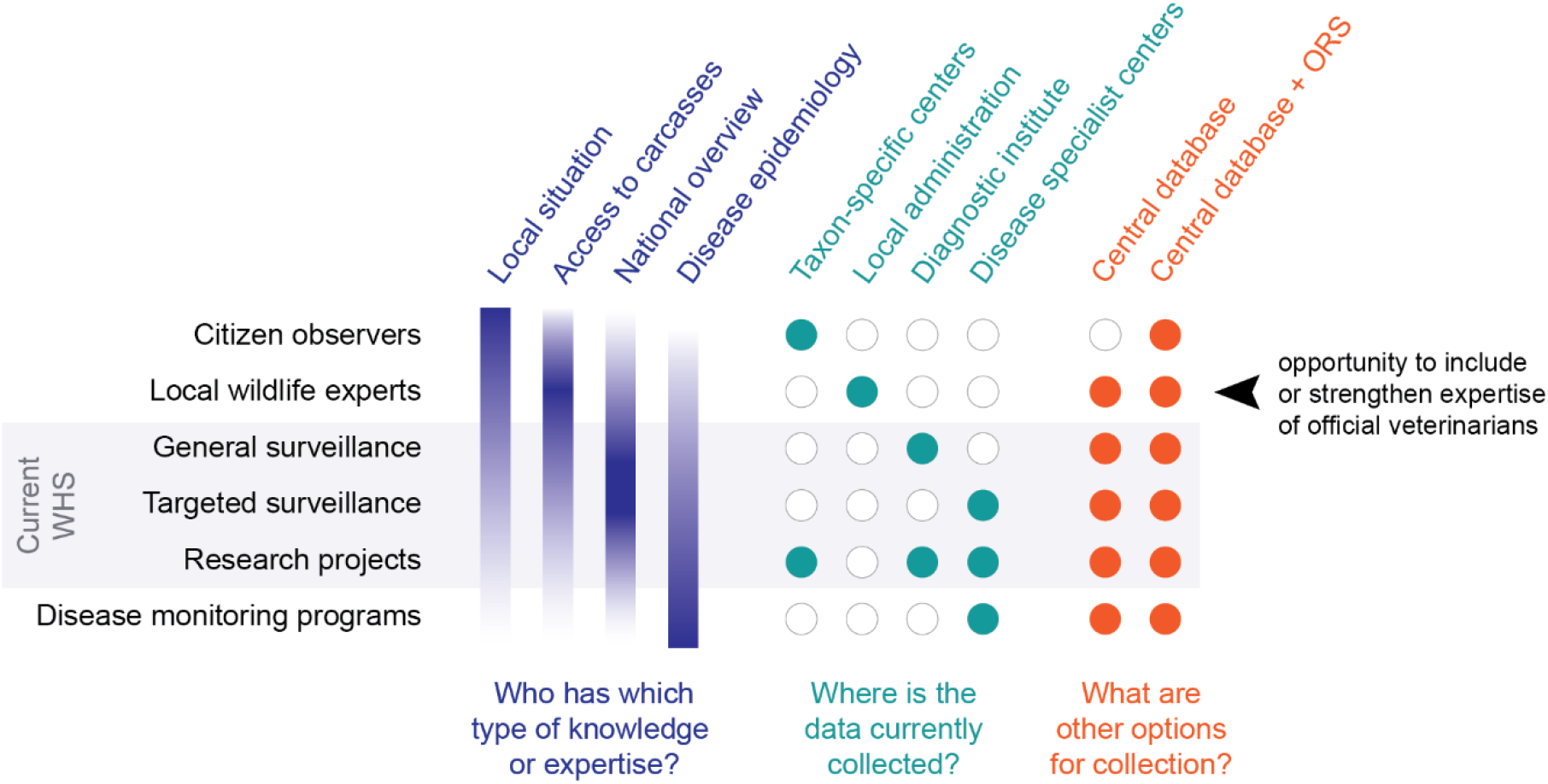
Knowledge and data sources, collection points, gaps, and recommendations. Wildlife health surveillance (WHS) knowledge and expertise is scattered across various levels of geographic areas and expertise. Citizens, local experts, general surveillance, targeted surveillance, research projects, and specialist centers each contribute different types of expertise (violet), and their knowledge is currently collected at different sites (teal). More holistic collection options would include a central database that is accessible for all contributors, and the inclusion of online reporting systems (ORS) in the data collection process (orange). Importantly, local veterinarians are presently not allowed to handle / treat wildlife except for euthanasia to end suffering, and official veterinarians are usually not trained to recognize or deal with wildlife disease.

#### A variety of measures could address the apparent information needs and information gaps identified in this study

Some of them are straightforward to implement – others require a concerted effort from a multitude of partners. **First**, current annual and quarterly reports which summarize relevant findings in the submitted cases issued by the diagnostic institute retrospectively at regular intervals are currently available only to a limited set of recipients in the federal offices and cantonal administration. Provided that data protection issues can be solved satisfactorily, these reports could be additionally distributed to rehabilitation centers and other non-cantonal institutions or made freely available on an online repository. **Second**, fragmented and federalist systems like the one presented absolutely require a central, consistent and accessible collection of numbers of animals found dead or diseased in a central database (Fig 5). Otherwise, efforts of multiple actors at multiple levels go to waste. Recently, the federal authorities have launched a project to incorporate more health-related data in the hunting statistics. A key part of such a central data collection is to include not just the resources for the database and the information management, but also for regular, real-time, modeling-based, epidemiological evaluation of the data to understand remaining blind spots and opportunities as well as needs for targeted disease surveillance. **Third**, the potential of the GWHS in guiding case selection and in reporting data could be expanded or strengthened (Fig 5). So far, field diagnoses have rarely been reported to the GWHS, resulting in information gaps in national records and therefore in reports to the World Organization of Animal Health and knowledge gaps in general. An inclusion of local veterinary experts would create additional needs with regard to training and education, a need that the GWHS host institution is able to expertly fill thanks to its academic setting. **Fourth**, online reporting systems (ORS) represent an option to harness decentrally available information, and thus fill wildlife health information gaps (Fig 5). A few countries have implemented ORS, for example based on mobile technologies [61]. Mobile-phone based reporting systems provide almost real-time data, which proved practicable especially in lower-resource or remote rural settings [62]. A study using an internet-based surveillance system to monitor avian influenza activities Twitter showed that such a systems might complement traditional surveillance systems and thus contribute to early detection of disease outbreaks [63]. Therefore, the establishment of an ORS for wildlife diseases and mortality in Switzerland has great potential to improve the knowledge of wildlife health across the country and to acknowledge the value of the collected data. If compatible with wildlife population monitoring systems or linked to other institutions, it could even generate synergies between wildlife disease surveillance and wildlife population monitoring, leading to so-called integrated monitoring [9,11] with the possibility of generating combined cartographic data. It would be desirable for such an ORS not to be limited to selected diseases, but to allow for reporting clinical signs and visible lesions (the base of the so called “syndromic surveillance”) as well as tentative (field) diagnoses for all animals found dead [64,65]. According to our results, most hunting authorities would support the introduction of an ORS. However, a project to introduce a nationwide uniform ORS would have to take existing concerns (additional effort, misdiagnosis, technical challenges) seriously and address them proactively and in a sociologically sound manner. At the same time, the information technology challenges (links to different databases, targeted access to certain data categories) would have to be professionally addressed by the IT side. Finally, it would have to be clarified in a legally secure manner where such a database can be located, who undertakes systematic evaluations and who gets access to the data. **Finally**, citizen observers can play a key role in wildlife health surveillance. The role of citizen science for targeted or general surveillance of wildlife disease including syndromic approaches has been explored previously [50]. Several citizen science reporting systems exist in Switzerland, mostly targeted at reporting native or invasive plants (www.infoflora.ch), bird sightings (www.ornitho.ch), but also general species occurrences (www.infofauna.ch). ORS could potentially incorporate diseases or syndromes, and feed into a central WHS database (Fig 5).

ORS based on field diagnoses carry the risk of missed diagnoses, i.e., diseases that are not recognized in the field because they lack clear visible signs and of misdiagnoses, i.e., the visible signs cannot clearly be assigned to one disease (e.g., central nervous symptoms). Therefore, the deployment of an ORS must include a) information of the submitter’s expertise and experience, b) some kind of measure of certainty (either added in the field or post-hoc based on submitter role, and c) an evaluation of how important it is in a given epidemiological setting to get each and every field diagnosis right. For some diseases, an ORS may allow to gather information where there was none at all before, which can be very valuable despite an inevitable certain level of mistakes. A risk-benefit analysis should therefore be part of ORS implementation and evaluation.

## Conclusions and Recommendations

For stakeholders and administration facing the challenge of building, improving or adapting a general wildlife health surveillance program, we offer the following recommendations. **Firstly**, changes in disease dynamics, in education and awareness, in focus species, and in population sizes need to be accounted for when projecting the capacities and resources of a GWHS. In particular, increasing population sizes of protected species can present unexpected challenges to a GWHS if their health is monitored in-depth. In a multi-stakeholder scenario, communicable, clear criteria might facilitate case selection and communication of caseload developments. Therefore, we suggest the development of careful case definitions and a decision tree that is tailored to the goals of a general wildlife health surveillance program, to the expectations of the mandating national authorities, and the needs of field partners. We also recommend engaging in regular open exchange to spot and discuss changing settings early on. **Secondly**, we recommend keeping an eye out for information gaps in common or easily recognizable diseases, species that are not hunted, or regions that are most distant to the monitoring institute, and to implement annual or bi-annual focus species programs to cover any systematic gaps through targeted surveillance programs. **Thirdly**, we recommend the systematic implementation of data sharing processes between all institutions involved in aspects of wildlife health monitoring which should be independent from personel connections. Also, considering the growing role of citizen observers in environmental research, and the data gap for garden wildlife in a hunting-heavy surveillance system, the inclusion of citizen observations seems advisable. For both issues, an ORS represents an option to harness decentrally available information, and thus fill wildlife health information gaps.

## Acknowledgements

We thank all cantonal hunting authorities (including game wardens, hunters, hunting wardens) for participating in the online survey as well as the hunting inspectors for taking time for the telephone interviews. We also thank Adrian Arquint, Thomas Gerner, Daniela Hadorn, Barbara Thür and Cordia Wunderwald for providing valuable inputs during conceptualization and progress of the study. Finally, we thank Robine Schoch for her help in the retrospective case analysis.

## Supporting information

*During the pre-print stage, supporting information is available from the authors on request*.

S 1 Appendix. Survey questions and answers

S 2 Appendix. Variables

S 3 Appendix. Questions needs assessment of hunting administrations

S 4 Appendix. Questions needs assessment of non-cantonal institutions

S 5 Appendix. Overview of univariable analysis

S 6 Appendix. Overview of multivariable analysis

S 7 Appendix. Decision tree

## Author contributions

**Conceptualisation:** Marie-Pierre Ryser-Degiorgis, Saskia Keller, Irene Adrian-Kalchhauser

**Data Collection:** Elisabeth Heiderich, Saskia Keller, Francesco Origgi, Samoa Zürcher-Giovannini, Stéphanie Borel, Iris Marti, Patrick Scherrer, Simone Roberto Rolando Pisano

**Data Curation:** Elisabeth Heiderich, Brian Friker

**Formal Analysis:** Elisabeth Heiderich, Brian Friker

**Funding Acquisition:** Marie-Pierre Ryser-Degiorgis, Irene Adrian-Kalchhauser

**Methodology:** Marie-Pierre Ryser-Degiorgis

**Project Administration:** Marie-Pierre Ryser-Degiorgis, Saskia Keller, Irene Adrian-Kalchhauser

**Supervision:** Marie-Pierre Ryser-Degiorgis, Irene Adrian-Kalchhauser

**Visualization:** Elisabeth Heiderich, Irene Adrian-Kalchhauser

**Writing – Original Draft Preparation:** Elisabeth Heiderich

**Writing – Review & Editing:** Marie-Pierre Ryser-Degiorgis, Mirjam Pewsner, Saskia Keller, Irene Adrian-Kalchhauser, Brian Friker, Francesco Origgi, Samoa Zürcher-Giovannini, Stéphanie Borel, Iris Marti, Patrick Scherrer, Simone Roberto Rolando Pisano

## Notes

### Competing Interest Statement

The authors have declared no competing interest.

